# A probabilistic approach to explore signal execution mechanisms with limited experimental data

**DOI:** 10.1101/732396

**Authors:** Michael A. Kochen, Carlos F. Lopez

**Affiliations:** Department of Biomedical Informatics, Vanderbilt University, Nashville, TN; Department of Biochemistry, Vanderbilt University, Nashville, TN

## Abstract

Mathematical models of biochemical reaction networks are central to the study of dynamic cellular processes and hypothesis generation that informs experimentation and validation. Unfortunately, model parameters are often not available and sparse experimental data leads to challenges in model calibration and parameter estimation. This can in turn lead to unreliable mechanistic interpretations of experimental data and the generation of poorly conceived hypotheses for experimental validation. To address this challenge, we evaluate whether a Bayesian-inspired probability-based approach, that incorporates available information regarding reaction network topology and parameters, can be used to qualitatively explore hypothetical biochemical network execution mechanisms in the context of limited available data. We test our approach on a model of extrinsic apoptosis execution to identify preferred signal execution modes across varying conditions. Apoptosis signal processing can take place either through a mitochondria independent (Type I) mode or a mitochondria dependent (Type II) mode. We first show that *in silico* knockouts, represented by model subnetworks, successfully identify the most likely execution mode for specific concentrations of key molecular regulators. We then show that changes in molecular regulator concentrations alter the overall reaction flux through the network by shifting the primary route of signal flow between the direct caspase and mitochondrial pathways. Our work thus demonstrates that probabilistic approaches can be used to explore the qualitative dynamic behavior of model biochemical systems even with missing or sparse data.

## Introduction

The complex dynamics of biochemical networks, stemming from numerous interactions and pathway crosstalk, render signal execution mechanisms difficult to characterize [1, 2, 3]. Mathematical modeling of biochemical networks has become a powerful compliment to experimentation for generating hypotheses regarding the underlying mechanisms that govern signal processing and suggesting targets for further experimental examination [4, 5]. Models of biochemical reaction networks, often based on a mass action kinetics formalism, are built to represent known pathway mechanics with knowledge garnered from years or even decades of experimentation [6, 7]. Although these models have yielded important predictions and insights about biochemical network processes, they also depend on kinetic rate parameters and protein concentrations that are often poorly characterized or simply unavailable. A typical workaround is to employ model calibration methods to estimate suitable parameter values via optimization to protein concentration time course data [8, 9, 10]. However, the data needed for parameter optimization is often scarce, leading to the possibility of multiple parameter sets that fit the model to that data equally well but exhibit different dynamics. [7, 9]. This poses a challenge for the study of dynamic network processes as the mode of signal execution can be highly dependent on a specific parameter set and could in turn lead to inadequate model-based interpretation. A computational approach that enables the exploration of biochemical signal execution mechanisms from a probabilistic perspective, constrained only by available data, would facilitate a rigorous exploration of network dynamics and accelerate the generation of testable mechanistic hypotheses [11].

In this work, we investigate whether a Bayesian-inspired probabilistic approach can identify network signal execution mechanisms in extrinsic apoptosis restricted only by experimental observations. Two execution phenotypes have been identified for extrinsic apoptosis signaling: a mitochondria independent (Type I) phenotype, whereby initiator caspases directly activate effector caspases and induce cell death, and a mitochondria dependent (Type II) phenotype whereby initiator caspases engage the Bcl-2 family of proteins, which ultimately lead to effector caspase activation (see Box 1 for biology details). Most mammalian cells execute apoptosis via the Type II mechanism, yet the Type I mechanism plays a central role in specific cell types, particularly certain types of lymphocytes [12]. A significant body of experimental and modeling work has identified key regulators for Type I vs Type II execution (see Box 1). However, it is still unclear how network structure and the interplay among multiple regulators can modulate signal execution for either cell type. A more traditional approach would prescribe intricate and detailed experimental measurements of cellular response to yield the desired data and improve our understanding of signal execution. However, the time and cost associated with such experiments makes it unlikely, and at times infeasible, to obtain said data. It is here that we see probabilistic inference approaches as complementary to experimentation, providing qualitative insights about signal execution mechanisms by integrating the expected parameter space subject only to available computer time. Here we demonstrate that a probabilistic approach, constrained by network structure or molecular concentrations, can identify the dominant signal execution modes in a reaction network. Specifically, we demonstrate the dependence of Type I or a Type II cellular apoptosis execution on network structure and chemical-species concentrations. We use expected values for quantifiable in silico experimental outcomes as metrics for comparisons of signal flow through different pathways of the network and subnetworks in order to identify how regulators affect execution modes. We introduce two complementary approaches that can be used in tandem to explore signal execution modulation. We first define a *multimodel exploration method* to explore multiple hypothesis about apoptosis execution by deconstructing an established apoptosis network model into functional subnetworks that effectively represent in silico knockout experiments. We also define a *pathway flux method* to characterize the signal flux through specific network pathways within the chosen canonical network. Combined, these two approaches enable us to qualitatively identify key network components and molecular regulator combinations that yield mechanistic insights about apoptosis execution. Our approach is generalizable to other mass action kinetics-based networks where signal execution modes play important roles in cellular outcomes. This work leverages Nested Sampling algorithm methods to efficiently calculate expected values on high performance computing (HPC) platforms, both of which are seldom used in biological applications. In this manner we are able to carry out the necessary calculations to consider the entirety of the proposed parameter space and estimate expected values within the timespan of hours to days.

### Box 1. Extrinsic apoptosis execution

Extrinsic apoptosis is a receptor mediated process for programmed cell death. The Type I/II phenotypes for the extrinsic apoptosis system were first described by Scaffidi et al. [13]. In that work they examined several cell lines and classified them into those that required the mitochondrial pathway to achieve apoptosis (Type II) and those that don’t (Type I). They made several interesting conclusions. They found that Type II cells had relatively weak DISC formation, that both phenotypes responded equally well to receptor mediated cell death, that there was a delay in caspase activation in Type II cells, and that caspase activation happened upstream of mitochondrial activation in Type I cells and downstream in Type II. More recently, XIAP has also been put forth as a critical regulator in the choice of apoptotic phenotype. In Jost et al. [14] they examined hepatocytes (Type II cells) and lymphocytes (Type I cells) under different conditions to examine the role XIAP plays in Type I/II determination. They made several observations upon Fas ligand or Fas-antibody induced apoptosis such as higher levels of XIAP in Type II cells and higher caspase effector activity in XIAP/Bid deficient mice versus apoptosis resistant Bid-only knockouts. In all, they concluded that XIAP is the key regulator that determines the choice of pathway.

Extrinsic apoptosis is initiated when a death inducing member of the tumor necrosis factor (TNF) superfamily of receptors (FasR, TNFR1, etc.) is bound by its respective ligand (FasL, TNF-α, etc.), setting off a sequence biochemical events that result in the orderly deconstruction of the cell [15]. The first stage of this sequence is the assembly of the DISC at the cell membrane ① and the subsequent activation of Caspase-8. Upon ligand binding and oligomerization of a receptor such as FasR or TRAIL, an adapter protein, like FADD (Fas-associated protein with death domain), is recruited to the receptors cytoplasmic tail [16, 17, 18]. FADD, in turn, recruits Caspase-8 via their respective death effector domains (DEDs), thus completing DISC formation [17, 18]. Other DISC components could also be included here, such as the regulator cFlip [19]. Once recruited, proximal Procaspase-8 monomers dimerize, inducing their autoproteolytic activity and producing active Caspase-8 [20, 21, 22].

After Caspase-8 activation the apoptotic signal can progress down two distinct pathways that both lead to the activation of Caspase-3 and the ensuing proteolysis of downstream targets. One pathway consists of a caspase cascade in which active Caspase-8 directly cleaves and activates Caspase-3 ② [23], while another, more complex pathway is routed through the mitochondria. In the mitochondrial pathway Caspase-8 cleaves the pro-apoptotic Bcl-2 family protein Bid in the cytosol, which then migrates to the mitochondria ③ where it initiates mitochondrial outer membrane permeabilization (MOMP) and the release of pro-apoptotic factors that lead to Caspase-3 activation [24, 25].

MOMP has its own set of regulators that govern the strength of apoptotic signaling through the mitochondria ④. After Caspase-8 activated Bid, (tBid), migrates to the mitochondria it activates proteins in the outer mitochondrial membrane, such as Bax, that subsequently self-aggregate into membrane pores and allow exportation of Cytochrome-c and Smac/DIABLO to the cytosol [26]. Bid and Bax are examples of pro-apoptotic proteins from the Bcl-2 family, all of which share BH domain homology [27]. Other members of this family act as MOMP regulators; the anti-apoptotic Bcl-2, for example, binds and inhibits both Bid and Bax while the pro-apoptotic Bad similarly binds and inhibits its target, Bcl-2 [28, 29, 30, 31]. Many other pro- and anti-apoptotic members of the Bcl-2 family have been discovered and together regulate MOMP [32].

Regardless of which pathway is chosen, the intermediate results are Caspase-3 activation and subsequent cleavage of PARP ⑧, a proxy for cell death in the analyses here [33, 34]. XIAP (X-linked inhibitor of apoptosis protein) is an inhibitor of Caspase-3 and has been proposed to be a key regulator in determining the Type I/II apoptotic phenotype of a cell [35]. XIAP sequesters Caspase-3 but also contains a ubiquitin ligase domain that directly targets Caspase-3 for degradation. The inhibitor also sequesters and inhibits the Caspase-3 activating Caspase-9 residing within the apoptosome complex [36, 37, 38]. Apoptosome formation is initiated by Cytochrome-c exported from the mitochondria during MOMP ⑤. Cytochrome-c induces the protein APAF-1 to oligomerize and subsequently recruit and activate Caspase-9, thus forming the complex [39]. Another MOMP export, the protein Smac/DIABLO ⑥, binds and inhibits XIAP, working in tandem with Cytochrome-c to oppose XIAP and carry out the apoptosis inducing activity of the Type II pathway [40]. Finally, Procaspase/Caspase-6 constitutes a feed forward loop between Caspase-3 and Caspase-8 ⑦ [41].

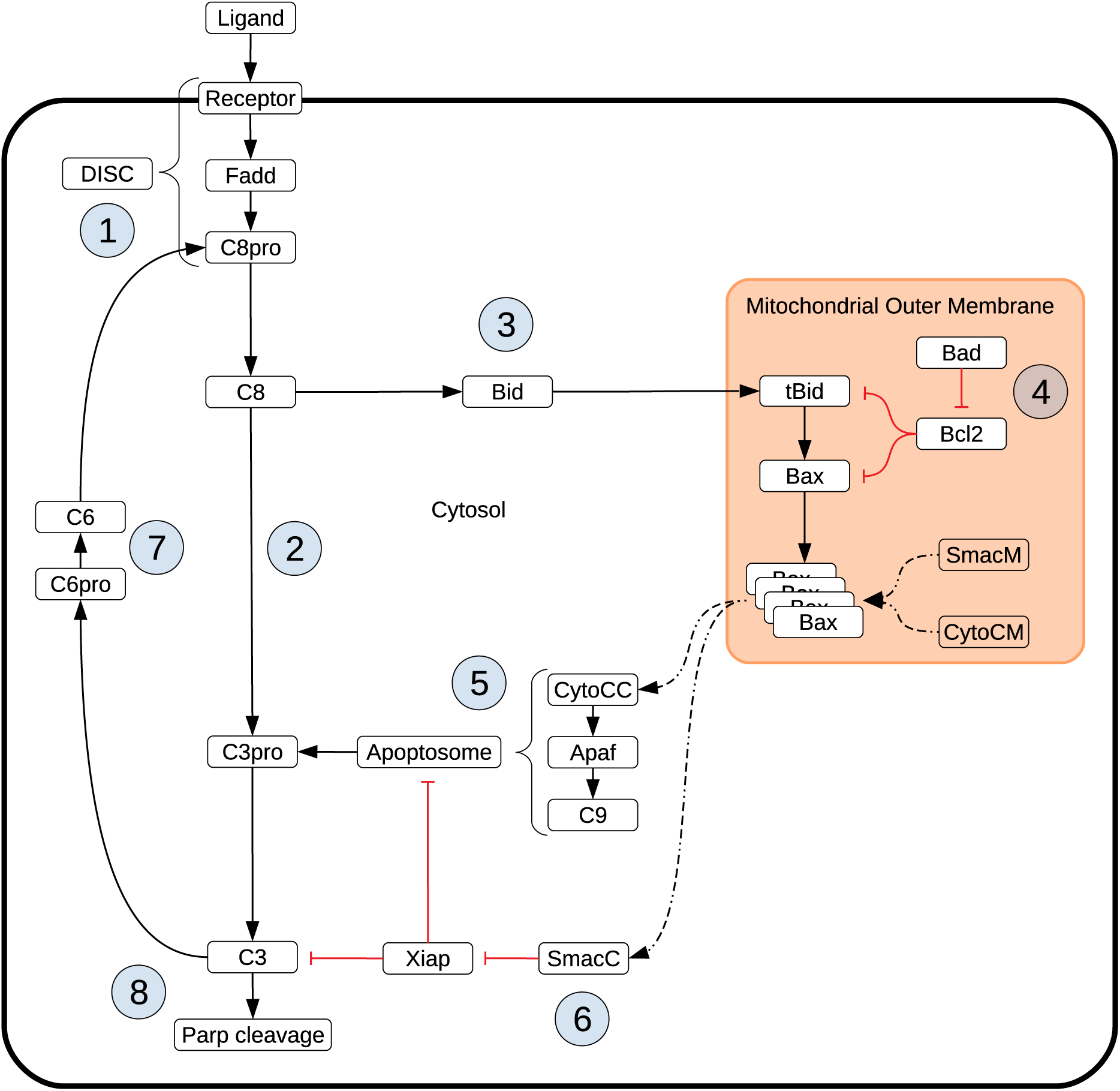
Box 1: Schematic of apoptotic signal flow through Type I and II pathways.

## Methods

### Apoptosis model and simulations

The base model used in this work is a modified version of the Extrinsic Apoptosis Reaction Model (EARM) from Lopez et al. (EARM v2.1) [7]. The original EARM was simplified to reduce complexity and lower the number of parameters, but still retains the key features of the network for apoptosis execution. Specifically, we reduced the molecular complexity of mitochondrial outer membrane permeabilization (MOMP) down to a representative set of Bcl-2 proteins that capture the behavior of activators, inhibitors, effectors, and sensitizers. We also eliminated intermediate states for Cytochrome c and Smac to streamline effector caspase activation, and we added an explicit FADD molecule, an adapter protein in the death-inducing signaling complex (DISC), to achieve a more realistic representation of signal initiation. Overall, EARM v2.1 is comprised of 16 chemical species at non-zero initial concentrations, 50 total chemical species, 62 reactions, and 62 kinetic parameters. The modified model was recalibrated to recapitulate the time-dependent concentration trajectories of truncated Bid, Smac release from the mitochondria, and cleaved PARP analogous to the approach reported previously [42] (Figure S1). The modified EARM, and all derivative models, were encoded in PySB. All simulations were run using the mass action kinetics formalism as a system of ordinary differential equations (ODEs) using the VODE integrator in SciPy within the PySB modeling framework. All data results, representative models, and software are distributed with open-source licensing and can be found in the GitHub repository https://github.com/LoLab-VU/BIND.

### Expected value estimation

The expected value for a quantifiable outcome is, by definition, the integral of an objective function that represents that outcome over the normalized distribution of parameters. This is analogous to the estimation of Bayesian evidence where a likelihood function is likewise integrated over a normalized distribution. We can thus use existing, established, Bayesian evidence estimation methods and software to estimate expected values by simply substituting the objective function for the likelihood function in the integral calculation. The remainder of this section and the next provide an overview of the evidence estimation methods and tools that we have repurposed for expected value calculations.

Bayesian evidence is the normalizing term in a Bayesian calculation and typically provides a measure for model comparison with regard to their fit to experimental data. It is expressed as:

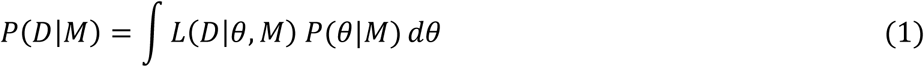

Where *M* is the model under consideration, *D* is the experimental data, θ is a specific set of parameter values, *L*(*D*|*θ, M*) is the likelihood function describing the fit of the data to the model under those parameter values, and *P*(*θ*|*M*) is the prior distribution of parameters. An efficient method for evidence calculation is nested sampling. This method simplifies the evidence calculation by introducing a prior mass element *dX* = *P*(*θ*|*M*) *dθ* that is estimated by (*X*_*i*−*i*_ − *X*_*i*_) where *X*_*i*_ = *e*^−*i/N*^, *i* is the current iteration of the algorithm, and *N* is the total number of live points. The evidence is then written as

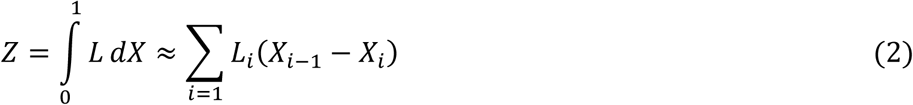

Initialization of the algorithm is carried out by randomly selecting an initial population of parameter sets (points in parameter space) from the prior distribution, scoring each one with the likelihood function, and ranking them from *L*_*high*_ to *L*_*low*_. At each iteration of the algorithm a new set of parameter values is selected and scored. If that score is higher than *L*_*low*_, then it is added to the population, at the appropriate rank, and *L*_*low*_ is removed from the population and added to the evidence sum (2).

### Nested sampling software

All expected value estimates in this work are calculated with MultiNest, a nested sampling-based algorithm designed for efficient evidence calculation on highly multimodel posterior distributions [44, 45]. MultiNest works by clustering the live points (population of parameter sets) and enclosing them in ellipsoids at each iteration. The enclosed space then constitutes a reduced space of admissible parameter sets. This lowers the probability of sampling from low likelihood areas and evaluating points that will only be discarded. The evidence estimate is accompanied by an estimate of the evidence error. The algorithm terminates when the presumed contribution of the highest likelihood member of the current set of live points, *L*_*high*_*X*_*i*_ is below a threshold. Here, we use a threshold of 0.0001 and a population size and 16,000 unless otherwise noted. The population size of 16,000 was found to be an acceptable compromise between precision and computational austerity for the model sizes and in silico experiments performed in this study. See [44, 45], for more details on the MultiNest algorithm. We use MultiNest with the Python wrapper PyMultiNest [46], which facilitates the integration with PySB into the parameter sampling pipeline.

### Multimodel exploration analysis

We carried out an analysis analogous to knockout experiments to investigate the contribution of different network components to the overall dynamics of the apoptosis execution network. We broke down the EARM network into six subnetworks and compared their likelihood of achieving apoptosis across increasing concentrations of the regulator XIAP. A standard proxy for apoptosis execution is cleavage of the protein PARP. We therefore define the proportion of cleaved PARP, relative to total PARP, as a metric for effective apoptosis execution. We defined the objective function that represents the amount of cleaved PARP as:

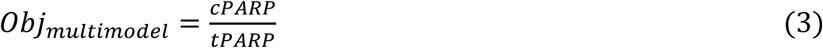

where *cPARP* is the amount of PARP that has been cleaved and *tPARP* is the total amount of PARP in the system. When this objective function is substituted into equation (1) in place of the likelihood function, we obtain the expected value, the average over the chosen prior parameter range, for the proportion of PARP that has been cleaved at the end of the in silico experimental simulation. We compare PARP cleavage for different subnetworks and regulatory conditions only in qualitative terms and as a *relative* measure of the expected outcome.

### Pathway flux analysis

We also explored the effect of molecular regulators of Type I vs Type II execution relative to the apoptosis signal flux through the network, as we have done in previous work [49]. Briefly, signal flux is defined as the chemical reaction flux in units of molecules per unit time, that traverses through a given pathway. In the apoptosis network there are two potential pathways that can lead to Caspase-3 activation and subsequently PARP cleavage. In the direct caspase pathway initiator caspases, represented here as “Caspase-8”, directly cleave and activate the effector caspases, represented here as “Caspase-3”. By contrast, in the mitochondrial pathway, effector caspases are activated via the apoptosome, and are dependent on MOMP. Therefore, the dominant pathway responsible for Caspase-3 activation defines the route of the signal. To estimate the flux through one of these pathways, we define the objective function as:

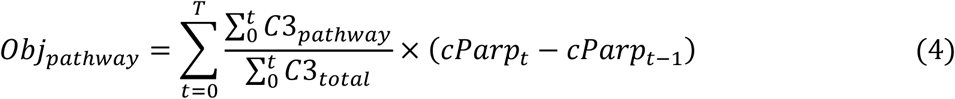

where *t* represents time in seconds, 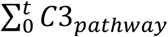 is the amount of Caspase-3 activated via the target pathway up to time *t*, 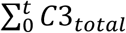 is the total Caspase-3 activated up to time *t*, and 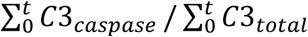 is the proportion of activated Caspase-3 that was produced via the target pathway up to time *t*. (*cParp*_*t*_ − *cParp*_*t*−1_) is the total PARP that has been cleaved, and activated, by Caspase-3 from time *t* − 1 to time *t*.Thus, at any given time *t* we can estimate the amount of Caspase-3 that has been activated through a specific pathway. Multiplication of these two terms returns an estimate for the amount of PARP cleaved via the specific pathway at time *t*. Summing over *T* then returns an estimate for the total apoptosis signal flowing through the target pathway. Like the PARP cleavage objective function, the signal flux objective substituted into equation (1) produces an estimate of the average flux over a defined prior distribution. We estimated this quantity over increasing concentrations of the molecular regulator XIAP, but also at high and low levels of the DISC components FADD and Caspase-8. The total signal flux was estimated by summing the flux estimate for both the direct caspase and mitochondrial pathways.

### Parameter ranges and initial conditions

The prior distribution takes the form of a set of parameter ranges, one for each reaction rate parameter. The ranges used here span four orders of magnitude around generic reaction rates deemed plausible [4] and are specific to the type of reaction taking place. The ranges of reaction rate parameters, in Log10 space, are 1^st^ order forward: [-4.0, 0.0], 2^nd^ order forward: [−8.0, −4.0], 1^st^ order reverse: [-4.0, 0.0], catalysis: [-1.0, 3.0]. These ranges were also used in the calibration of the base model. Where possible, initial conditions were either collected from the literature [50, 51] or taken from a previous model of extrinsic apoptosis [7, 52]. Because the baseline model was designed to concur with Type II apoptotic data (see above), literature derived initial conditions were based on Type II Jurkat or Hela cell lines (Table S1).

### Expected value ratios

Evidence estimates are often used to select between two competing models by calculating the Bayes factor (i.e. the ratio of their evidence values). This provides a measure of confidence for choosing one model over another. We can likewise use the ratios of expected values to gain additional insights into the dynamical relationship between network components. To facilitate construction of expected value ratios (EVR) with a continuous and symmetric range, we define them as:

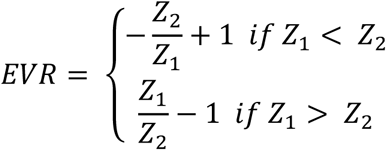

where Z_1_ and Z_2_ are the expected value estimates for two networks under comparison.

### Computational resources

Because of the high computational workload necessary for this analysis, a wide range of computational resources were used. The bulk of the work was done on the ACCRE cluster at Vanderbilt University which has more than 600 compute nodes running Intel Xeon processors and a Linux OS. As many as 300 evidence estimates were run in parallel on this system. Additional resources included two local servers, also running Intel processors and a Linux OS, as well as a small local four node cluster running Linux and AMD Ryzen 1700 processors. A detailed breakdown of CPU time can be found in the results section. In all, expected value estimates for 14 different networks/initial conditions were made across the range of XIAP concentrations. We estimate all 14 runs would take ∼9 days each on a typical university server with 32 cores/64 threads.

## Results

### Overview: A Bayesian-inspired approach to explore mechanistic hypotheses

Our overarching goal is to understand the mechanisms and dynamics of biochemical networks responsible for cellular commitment to fate, given incomplete or unavailable data. We take a probabilistic approach, similar to those used in Bayesian evidence-based model selection and multimodel inference, to compare model subnetworks and pathways with respect to apoptotic signal execution under various in silico experimental conditions and enable the generation of hypotheses regarding the underlying mechanisms of signal processing. Using this approach, we’ve employed two distinct but complimentary strategies.

The first is *Multimodel Exploration Analysis* (Figure 1, left path), wherein the network model is deconstructed into biologically relevant subnetworks and the probability of each subnetwork achieving apoptosis, under various regulatory conditions, is estimated via the calculation of an expected value for a quantifiable proxy of apoptosis. This differs from traditional model selection and multimodel inference applications where models are typically ranked based on their fit to experimental data and high-ranking models may be averaged to obtain a composite model [47, 48, 53, 54, 55, 56]. Here, we already have a model that captures key features of programmed cell death execution. Instead, we use the differences in expected values for a quantity that is representative of apoptosis to construct a composite picture of mechanistic evidence for apoptosis execution. To achieve this, we first tailor the objective function to represent signal execution strength, as measured by cleaved PARP concentration at the end of the simulation. The expected value derived from this objective function therefore describes the likelihood that the signal is effectively transmitted through a given network. It should be noted that Bayesian evidence, and by extension our expected value calculation, inherently incorporates model complexity as the objectives are integrated over normalized prior distributions [44, 53, 57]. As we will see, comparison of changes in signal strength through relevant subnetworks allows inferences to be made on the effect of the perturbed network regulator as well as various network components on the overall dynamics of the system. We focus primarily on understanding how Bayesian evidence for the caspase pathway compares to that of the complete network as these are most relevant for the analysis of Type I/II execution modes. This analysis will inform on how network components contribute to overall signal execution and provide mechanistic insights about the sensitivity of PARP cleavage to subnetwork components.

**Figure 1.**
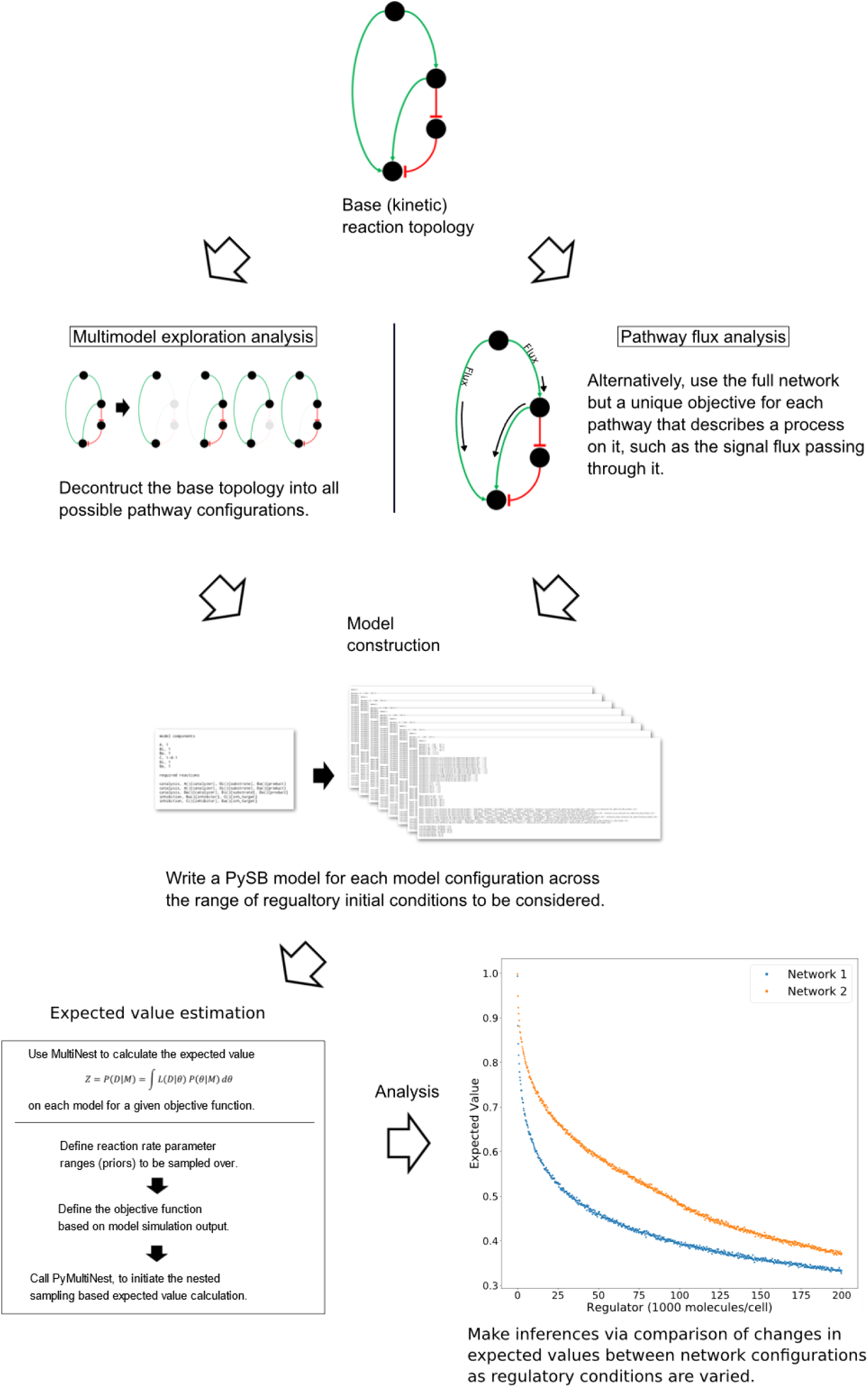
General workflow for the analysis of network dynamics using trends in expected values. The target network is first deconstructed into subnetworks that effectively represent in silico knockouts. A model for each subnetwork and each incremental set of regulatory conditions is then created and passed to an algorithm for estimation of the expected value for an aspect of signal transduction. The expected value is calculated via integration of a user-defined objective function that quantifies that aspect of signal transduction over a range of parameter values (the prior). The trends in the expected values over changing regulatory conditions are then compared to make qualitative inferences regarding network dynamics. In a complimentary method, the full model is retained but the objective function is targeted to different pathways. Inferences on network dynamics can again be made via comparison of the trends in the expected values.

The second strategy is *Pathway Flux Analysis* (Figure 1, right path), where we retain the complete network structure but instead tailor the objective functions to measure biochemical reaction flux through either the direct caspase or mitochondrial pathways. We primarily consider the influence of the apoptosis inhibitor XIAP on regulatory dynamics and phenotypic fate but also consider the regulatory effect of the death inducing signaling complex (DISC) and the anti-apoptotic protein Bcl-2, all of which have been found to be relevant to Type I vs Type II execution in different cell types [13, 14]. This analysis will inform on how molecular regulators modulate biochemical flux through the network and their influence on apoptosis completion as measured by PARP cleavage.

### Decomposition of the extrinsic apoptosis network and reductive analysis of the effects of XIAP

To investigate the effect of network substructures on apoptosis signaling, we build a composite description of system dynamics by observing variations in signal throughput, represented by expected values of PARP cleavage, between subnetworks (Figure 2A-F) relative to changes in regulatory conditions. We consider relative changes in expected PARP cleavage as the number of XIAP molecules is increased where a higher value indicates a stronger average signal over the prior range of parameter values. XIAP was varied from 0 to 200,000 molecules per cell in increments of 250 to explore how changes in XIAP affect the likelihood of apoptosis execution. For subnetworks that include the mitochondrial pathway, Bcl-2 (an anti-apoptotic protein) was eliminated, to explore Type I vs Type II activity independent of inhibitors that could confound signal throughput, and more closely simulate a cell that is “primed” for death [57]. All other initial values were fixed at the levels shown in supplementary Table S1. In the absence of XIAP all six subnetworks have PARP cleavage estimates greater than 0.98, (Figure 2 A: 0.993, B: 0.998, C: 0.992, D: 0.981, E: 0.998, F: 0.981, Table S2) indicating a robust apoptotic signal for each across the allowed range of parameters. The log-expected value version of Figure 2G along with estimated errors generated by MultiNest are displayed in Figure S2.

**Figure 2.**
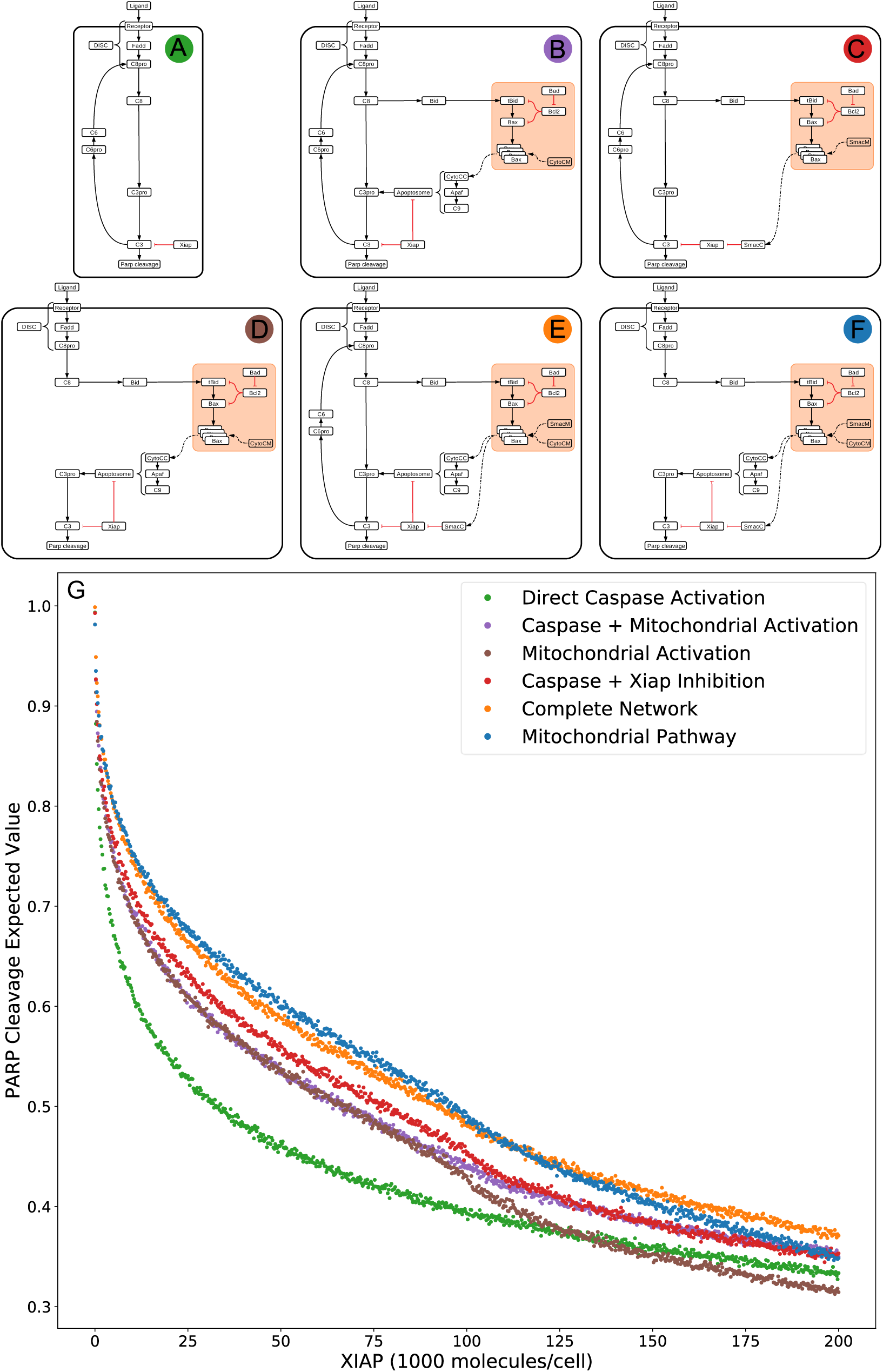
Extrinsic apoptosis subnetworks and the likelihood of achieving apoptosis. (A) The direct caspase subnetwork. (B) The direct caspase + mitochondrial activation subnetwork. (C) The direct caspase + mitochondrial inhibition of XIAP subnetwork network. (D) The mitochondrial activation subnetwork. (E) The complete network. (F) the mitochondrial subnetwork. (G) Trends in expected values for each of the networks in (A)-(F) over a range of values for the apoptosis inhibitor XIAP and for an objective function that computes the proportion of PARP cleavage (a proxy for cell death) at the end of the in silico experimental simulation.

The results in Jost et al. [14] imply that the cellular level of XIAP determines the preferred apoptosis pathway with higher levels specific to Type II cells and lower levels specific to Type I. To hypothesize a possible mechanistic explanation for this behavior we compared the expected PARP cleavage, over increasing concentrations of XIAP, for the direct caspase activation network against both the complete network and the isolated mitochondrial pathway network (Figures 2A and G green; 2E and G orange; 2F G blue respectively). This mimics reported experimental strategies to study Type I/II phenotypes and allows us to gauge the effect of XIAP on networks with and without a mitochondrial component [13, 35].

As XIAP levels increase we see differential effects on all subnetworks in the form of diverging expected value estimates, indicating differences in the efficacy of XIAP induced apoptotic inhibition. PARP cleavage values for the isolated caspase pathway (Figure 2G green) diverge from the complete network (Figure 2G orange) and mitochondrial pathway (Figure 2 blue) showing a steeper initial decline that diminishes as XIAP continues to increase. PARP cleavage values for the caspase pathway falls to 0.5 at an XIAP level of roughly 32,000. However, the complete network and mitochondrial pathways require XIAP levels nearly threefold higher with PARP cleavage reaching 0.5 at around 92,000 and 95,000 respectively.

Because the direct caspase activation pathway (Figures 2G green) is representative of the Type I phenotype, the disproportionate drop in its expected PARP cleavage as XIAP concentration increases is consistent with experimental evidence showing XIAP-induced transition from a Type I to a Type II execution mode [14]. The complete network, containing the full mitochondrial subnetwork, and mitochondrial only pathway are also affected by XIAP but exhibit resistance to its anti-apoptotic effects, a difference that is most prominent at moderate levels of the inhibitor. This suggests a dependence on mitochondrial amplification for effective apoptosis as XIAP increases from low to moderate levels. At higher levels of XIAP the PARP cleavage for the caspase pathway level off and the gaps between it and the two mitochondrial containing networks narrow. The disproportionate effect of XIAP inhibition of apoptosis on the caspase pathway suggests that the mechanism for XIAP induced transition to a Type II pathway can be attributed to differential inhibition of the apoptotic signal through the isolated caspase pathway vs a network with mitochondrial involvement.

The next two highest trends in expected values after that of the direct caspase network belong to the networks representing direct caspase activation plus mitochondrial activation and mitochondrial activation alone (Figures 2G purple and 2G brown). For most of the range with XIAP below 100,000 these two networks have largely overlapping PARP cleavage trajectories, despite the fact that the former has twice as many paths carrying the apoptotic signal. Near an XIAP level of 100,000 the two trends diverge as the decrease in PARP cleavage for the mitochondrial activation only network accelerates. This could be explained by XIAP overwhelming the apoptosome at these higher levels. The apoptosome is an apoptosis inducing complex (via Caspase-3 cleavage) consisting of Cytochrome c, APAF-1, and Caspase-9, and is an inhibitory target of XIAP. As XIAP increases past 125,000 the mitochondrial activation only PARP cleavage values fall below even the solo direct caspase values, possibly due to the two-pronged inhibitory action of XIAP at both the apoptosome and Caspase-3. An interesting observation here is that the addition of the direct caspase pathway to the mitochondrial activation pathway does not appear to increase the likelihood of achieving apoptosis for lower values of XIAP.

PARP cleavage values for the network representing direct caspase activation plus mitochondrial inhibition of XIAP are in red in Figure 2G. Below an XIAP level of 100,000 these values are consistently above the PARP cleavage values for the network representing direct caspase plus mitochondrial activation. Note that while direct caspase activation does not appear to increase the likelihood of achieving apoptosis when added to the mitochondrial activation pathway (Figure 2G purple) the amplification of the direct caspase activation via mitochondrial inhibition of XIAP leads to a higher likelihood than solo activation through the mitochondria. This suggests the possibility that the primary mechanism for mitochondrial apoptotic signal amplification, under some conditions, may be inhibition of XIAP, with direct signal transduction a secondary mechanism. Above an XIAP level of 100,000, the direct caspase with XIAP inhibition PARP cleavage values drop to levels roughly in line with the values for direct caspase activation plus mitochondrial activation, possibly due to the fact that Smac, the mitochondrial export that inhibits XIAP, is also set to 100,000 molecules per cell. Both, however, remain more likely to attain apoptosis than direct caspase activation alone.

The two subnetworks with the highest expected values for apoptotic signal execution are the complete network and the isolated mitochondrial pathway (Figures 2E orange and 2F blue). As previously mentioned, both of these networks contain the full mitochondrial pathway implying that this pathway supports resistance to XIAP inhibition of apoptosis. Between XIAP levels of 0 to 100,000 the two trends track very closely, with the mitochondrial only pathway showing a slight but consistent advantage for apoptosis execution. The average difference between an XIAP level of 20,000 and 80,000 is roughly 0.014, meaning we expect the average PARP cleavage to favor the mitochondrial only pathway by about 1.4 percentage points, which may seem unremarkable. Context matters however, and the context here is that the complete network has potentially twice the bandwidth for the apoptotic signal, namely the addition of the more direct caspase pathway. Together, this raises the possibility that under some conditions the caspase pathway is not a pathway but a sink for the apoptotic signal. In such a scenario, the signal through the caspase pathway would get lost as Caspase-3 is degraded by XIAP. Not until the signal through the mitochondrial pathway begins inhibiting XIAP could the signal proceed. Around the 100,000 level of XIAP the PARP cleavage trend for the mitochondrial pathway crosses below that for the complete network. This could be due to the parity with Smac, components of the apoptosome, or a combination of the two.

### Apoptosis signal strength dictates the signal route through the network

The results in Scaffidi et al. [13] indicate a strong phenotypic dependence on the strength of the apoptosis signal. Here we examine hypotheses made in that work and the interplay between the DISC and XIAP regulatory axes. We again increase XIAP from 0 to 200,000 molecules in increments of 250, but this time at a low number of DISC complexes by lowering the initial values of both the scaffold protein FADD and the initiator Caspase-8, from 130,000 to 100 molecules per cell. In addition to the *Multimodel Exploration Analysis* approach used in the previous section, we also use the *Pathway Flux Analysis* approach using the signal flux objective function (see Methods). In this way we attain a holistic view of network dynamics that incorporates both network structure and signal flux crosstalk from all possible pathways. Additional analysis of caspase and mitochondrial pathway signal flux over a range of values for both XIAP and Bcl-2 is displayed in Figure S3 and interpreted in Text S1.

Figure 3A displays the PARP cleavage expected values along with their low DISC counterparts. Two things are immediately apparent. PARP cleavage for the caspase pathway with a low number of DISC molecular components is lower across the entire range of XIAP concentrations. The complete network, on the other hand, shows almost no difference under low DISC conditions at lower values of XIAP. This supports the hypothesis that mitochondrial involvement is necessary to overcome weak DISC formation and that weak signal initiation constitutes a Type II trait [13].

**Figure 3.**
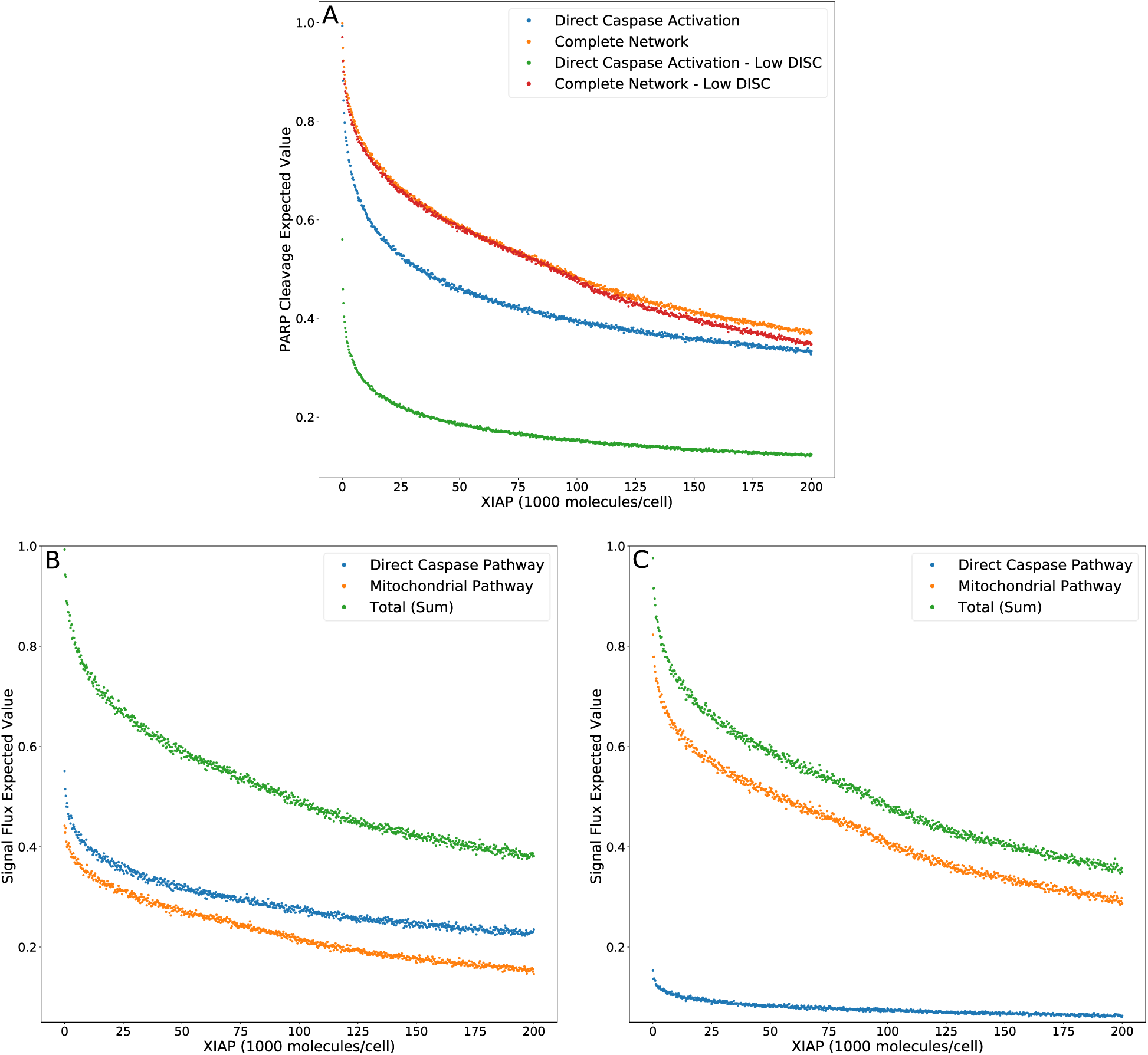
Expected values for PARP cleavage and pathway flux at low and high DISC component values. (A) Expected values for PARP cleavage for the caspase pathway and complete network under both low and high DISC conditions (100 and 130,000 molecules per cell of FADD and Caspase-8 respectively) over a range of XIAP values. (B) Expected values for signal flux through both pathways as well as the total signal flux under high DISC conditions. (C) Expected values for signal flux through both pathways as well as the total signal flux under low DISC conditions.

Figures 3B and 3C show expected values for signal flux through the caspase pathway and complete network, for high and low numbers of DISC components, respectively. At higher DISC values, signal flux through the caspase pathway is consistently higher than the flux through the mitochondrial pathway. At lower DISC values the signal flux through the mitochondrial pathway exceeds the flux through the caspase pathway. These results shed interesting mechanistic observations in the context of a previously proposed hypothesis stating that mitochondrial activation is downstream of Caspase-8 activation in Type I cells and upstream in Type II cells. If a weaker initial apoptosis cue does indeed push the signal through the mitochondrial pathway the initial activation of Caspase-8 would be weak and the amplifying activity of the mitochondria would ramp up the signal before Caspase-8 could directly activate Caspase-3. On the other hand, strong initial activation that pushes the signal through the caspase pathway would activate both Caspase-8 and Caspase-3 before MOMP becomes fully active. Also notable is the nearly identical trajectories of the total signal flux through the low and high DISC models. The average difference over the range of XIAP was only 0.011 (Table S3). This is consistent with observations that both Type I and Type II cells respond equally well to receptor mediated apoptosis [13].

Overall these results set up three mechanistic explanations for apoptosis execution. On one end, strong signal initiation and low XIAP results in the independence of apoptosis from the mitochondrial pathway. This behavior is consistent with Type I cells like the SKW6.4 cell lines [13]. Under this scenario our results imply that the majority of the signal flux is carried through the caspase pathway and we hypothesize that control of apoptosis is dominated by that pathway. On the other end of the spectrum weak signal initiation and moderate to high levels of XIAP result in a dependence on the mitochondrial pathway. Such behavior is consistent with Type II cells like Jurkat [13]. In this case our results strongly indicate that the majority of signal flux is carried through the mitochondrial pathway and we hypothesize that apoptosis execution is dominated by that pathway. In between these two extremes is the case with strong signal initiation, and moderate to high levels of XIAP levels with apoptotic dependence on mitochondrial activity. Such a scenario that is consistent with MCF-7 cell that are known to have traits of both phenotypes [13]. In this case, we found that the majority of the apoptotic signal is carried through the caspase pathway despite the dependence on the mitochondria and we hypothesize that the mitochondrial pathway acts to allow the apoptotic signal through the caspase pathway.

### Expected value ratios and XIAP influence on Type I/II apoptosis phenotype

Model selection methods typically calculate the evidence ratios, or Bayes factors, to choose a preferred model and estimate the confidence of that choice [59, 60].When comparing changes in likelihood of an outcome as regulatory conditions are altered we can similarly use ratios of expected values to provide additional information about evolving network dynamics under regulatory perturbations. To characterize the effect of XIAP on the choice of Type I or II apoptotic phenotype we calculated the expected value ratios (Figure 4A), for each value of XIAP between the caspase pathway and both the complete network and mitochondrial pathway. In these calculations, the denominator represents the caspase pathway so that higher values favor a need for mitochondrial involvement. An interesting feature of both the complete and mitochondrial expected value ratios is the peak and reversal at a moderate level XIAP (Figure 4B). This reflects the initially successful inhibition of the caspase pathway that decelerates relatively quickly as XIAP increases, and a steadier rate of increased inhibition on networks that incorporate the mitochondrial pathway. The ratios peak between 45,000 and 50,000 molecules of XIAP (more than double the value of its target molecule Caspase-3 at 21,000) and represent the optimal level of XIAP for the requirement of the mitochondrial pathway and attainment of a Type II execution. Given the near monotonic decline of the expected values for both pathways, representing increasing suppression of apoptosis, the peak and decline in the expected value ratios could represent a shift toward complete apoptotic resistance. Our results therefore complement the observations in Aldridge et al. where a similar outcome was observed experimentally [52].

**Figure 4.**
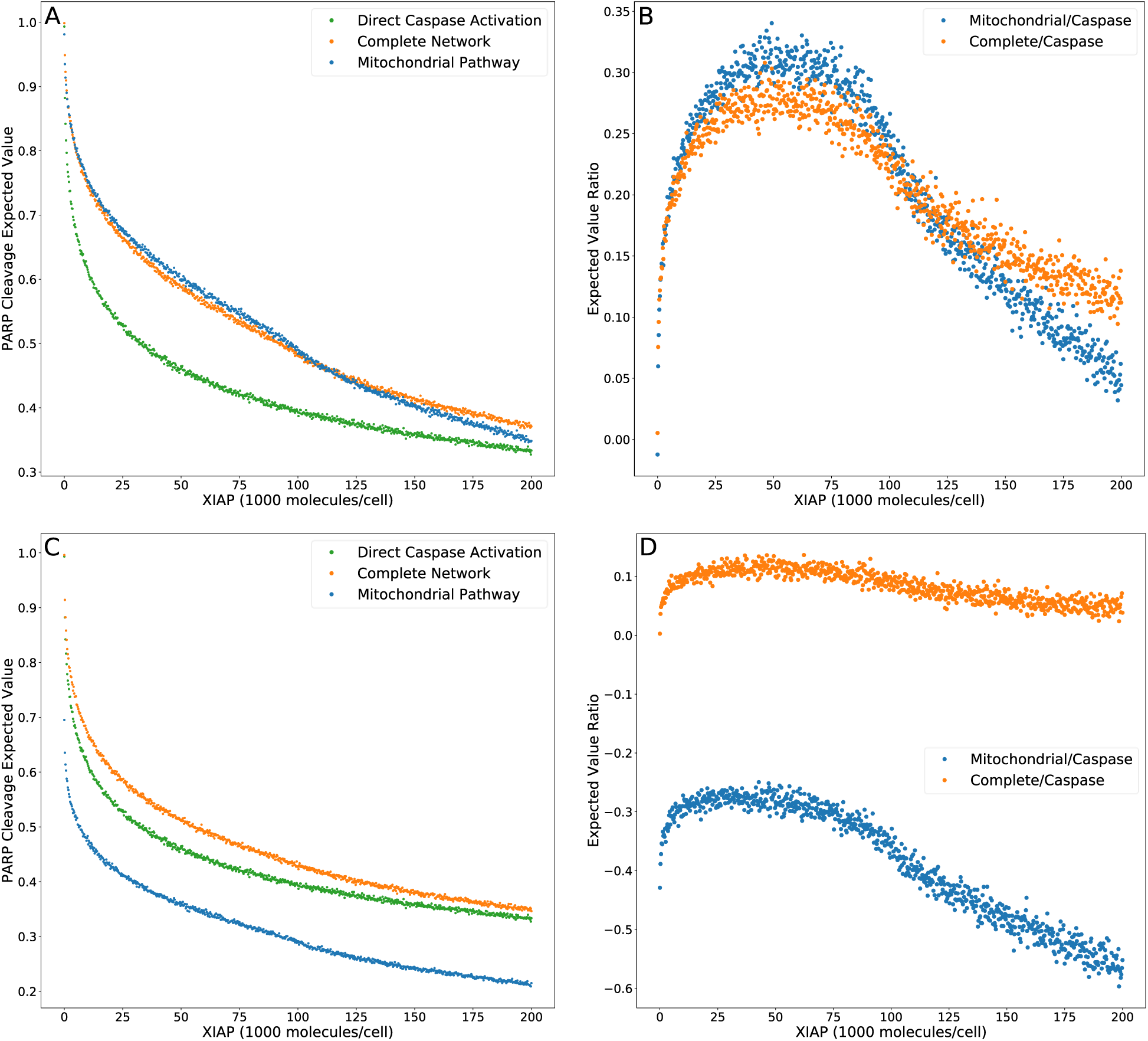
Trends in expected value ratios under increasing levels of the apoptotic inhibitor XIAP for an inhibited and uninhibited mitochondrial pathway. (A) Expected value trends for the caspase pathway (green), mitochondrial pathway (blue), and complete network (orange) with no MOMP inhibition. (B) Trends for the mitochondria/caspase (blue) and the complete/caspase (orange) expected value ratios from the trends in (A). (C) Expected value trends for the caspase pathway (green), mitochondrial pathway (blue), and complete network (orange) with MOMP inhibitory protein BCL-2 at 328,000 mol. per cell. (D) Trends for the mitochondria/caspase (blue) and the complete/caspase (orange) evidence ratios from the trends in (C).

A common technique to study apoptosis is to knockdown Bid, overexpress Bcl-2, or otherwise shut down MOMP induced apoptosis through mitochondrial regulation. This strategy was used in Jost et al. [14] to study the role of XIAP in apoptosis and in the work of Aldridge et al. to explore Type I vs Type II execution in different cell lines [60]. Taking a similar approach, we set Bcl-2 levels to 328,000 molecules per cell, in line with experimental findings [47], to suppress MOMP activity and recalculated the PARP cleavage expected values and their ratios (Figures 4C and 4D, Table S5). Under these conditions PARP cleavage for the mitochondrial pathway drop well below that of the direct caspase pathway, which is reflected in the expected value ratios trend as a shift into negative territory and indicate that the caspase pathway is favored. PARP cleavage for the complete network under MOMP inhibition is shifted closer to that for the caspase pathway at higher concentrations of XIAP but is still higher throughout the full range of XIAP. The peak in the associated expected value ratios is flattened as the level of XIAP increases from low levels, suggesting that increasing XIAP is less likely to induce a transition to a Type II phenotype in a system with an already hampered mitochondrial pathway. We note that complete inhibition of MOMP would result in uninformative mitochondrial pathway results. PARP cleavage expected values for the complete network would be indistinguishable from those for the direct caspase pathway and the complete/caspase ratios would simply flatline. However, our analysis shows that isolation of active biologically relevant subnetworks and direct comparison under changing molecular regulatory conditions, using trends in expected values, enables the extraction of information regarding pathway interactions and differential network dynamics.

### Precision vs computational cost

Increasing the precision of the expected value estimates and tightening their trendlines, is accomplished by increasing the number of live points in the nested sampling algorithm. The trade-off is an increase in the number of evaluations required to reach the termination of the algorithm and an accompanying increase in total computation time. Figures 5A and 5B display the required number of evaluations for the direct caspase and complete network at population sizes of 500, 1000, 2000, 4000, 8000, and 16,000, when run with the PARP cleavage objective function. For both models the number of evaluations roughly doubles for every doubling in population size. Figures 5C and 5D are the average estimated errors calculated by the MultiNest algorithm over each population size for the direct caspase and complete networks respectively. As expected, error estimates fall roughly as *n*^−1*/*2^ [61], signifying clear diminishing returns as the number of live points is increased. The average CPU process times, as estimated by Python’s `time.clock()` method, are given in Figures 5E and 5F for the direct caspase and complete networks respectively. Despite the greater number of required evaluations for the direct caspase network the average clock times for the complete network is significantly higher. At a population of 16,000 the caspase network had an average clock time of 11,964 seconds compared to 76,981 for the complete network. Data for figure 5 can be found in Table S6.

**Figure 5.**
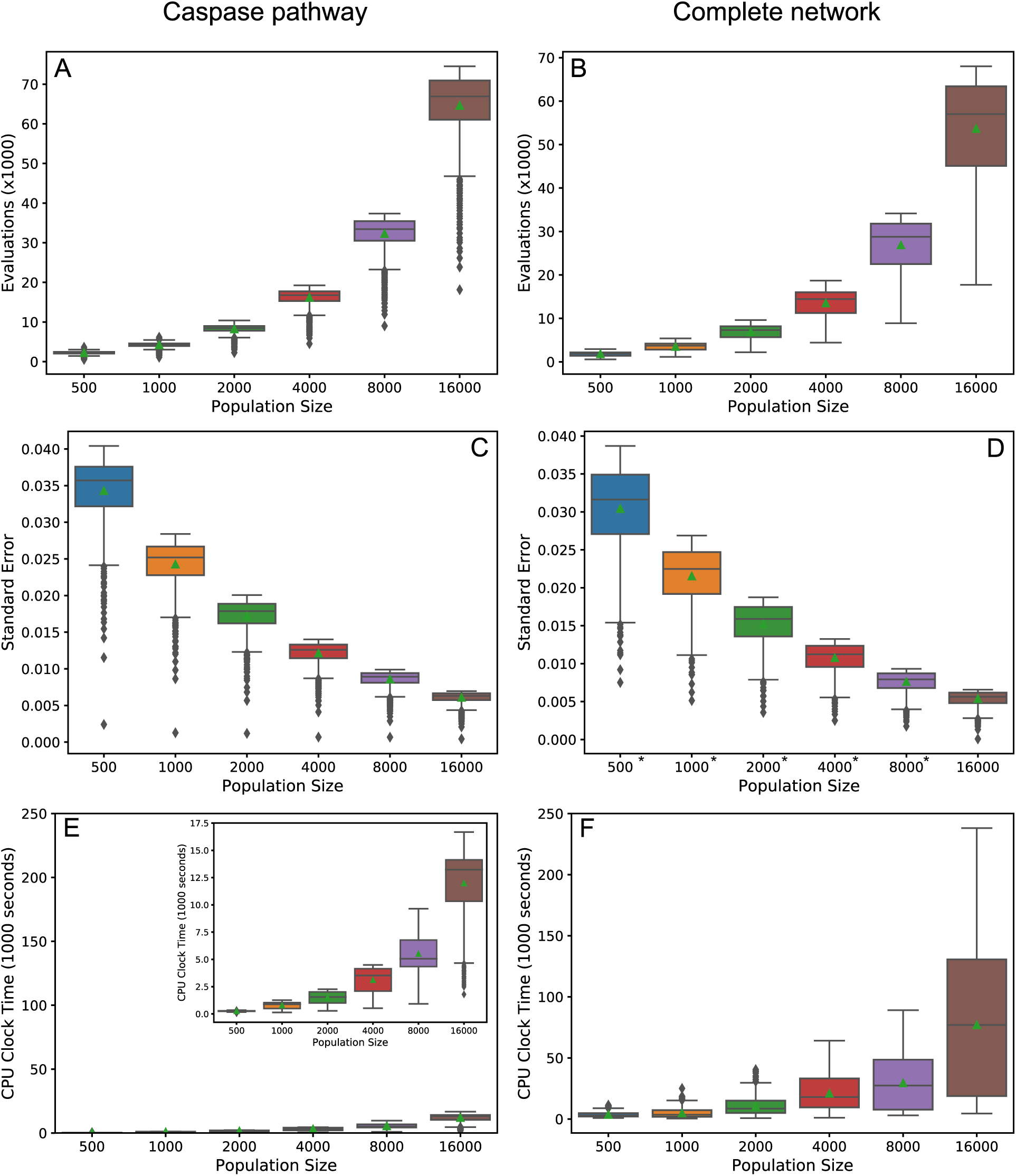
Precision vs. computational cost. (A) and (B) Average number of evaluations before termination of the MultiNest algorithm over a range of population sizes for the caspase pathway and complete network respectively. (C) and (D) Average of error estimates from MultiNest for each population size and the caspase and complete networks. (E) and (F) Average estimated CPU clock time over each population size for the caspase and complete networks respectively. *MultiNest was unable to estimate the error at XIAP = 0.

Ultimately, the choice of population size for the methods we have laid out here will depend on the networks to be compared, the objective function, and how well the trends in the expected values must be resolved in order to make inferences about network dynamics. For example, at a population size of 500 the trend in the PARP cleavage expected values for the direct caspase pathway is clearly discernable from that for the mitochondrial pathway and the complete network, but the latter two are largely overlapping (Figure S4A). At higher population levels, however, two distinct mitochondrial and complete PARP cleavage trends become apparent (Figure S4K). If expected value ratio trends are desired then the choice of population size must take into consideration the amplification of the noise from both expected value estimates (see Figures S4(B, D, F, H, J, L) for complete/caspase PARP cleavage expected value trends).

## Discussion

Characterizing information flow in biological networks, the interactions between various pathways or network components, and shifts in phenotype upon regulatory perturbations is a standing challenge in molecular biology. Although comparative analysis of signal flow within a network is possible with current computational methods, the dependence of physicochemical models on unknown parameters makes the computational examination of each network component highly dependent on costly experimentation.

To take advantage of the enormous amount of existing knowledge encoded in these physicochemical networks without the dependence on explicit parameter values we take a probabilistic approach to the exploration of changes in network dynamics. By integrating an objective function that represents a simulated outcome over parameter distributions derived from existing data we obtain the likelihood of attaining that outcome given the available information about the signaling pathways. The qualitative exploration of network behavior for various in silico experimental setups and regulatory conditions is then attainable without explicit knowledge of the parameter values. We demonstrate the utility of the method when applied to the regulation of extrinsic apoptosis. Networks that incorporate an active mitochondrial pathway displayed a higher resistance to apoptotic inhibition from increasing levels of XIAP, consistent with experimental evidence that XIAP induces a Type II phenotype [14]. Also in line with experimental evidence [13] are the results that suggest low/high signal initiation is consistent with Type II/I phenotype respectively and that both types achieve apoptosis equally well.

A potential limitation of this probabilistic approach to the study network dynamics is the computational cost. A number of factors affect the run time of the algorithm including the size of the model, the objective function, and the desired precision. Fortunately, reducing the resolution (the number of in silico experiments for which an expected value is estimated) and the precision (the population size) can drastically reduce the cost and in many cases the method will still be viable. One aspect of the method that is severely restrictive is the number of model components that can be varied in the same run since the computational cost increases exponentially with each additional variable. Reasonable parameter distributions must also be chosen, preferably based on existing data. Here we were able to use generic but biologically plausible ranges with uniform distributions to produce results that were qualitatively consistent with previous experimental results. These in silico generated qualitative results allow us to make mechanistic hypotheses from existing data over a period of weeks rather than the months or years that would be required to attain this information with experimental approaches. Our results therefore support probabilistic approaches as a suitable complement to experimentation and a shift from purely deterministic models with a single optimum parameter set to a probabilistic understanding of mechanistic models of cellular processes.

## Conclusions

In this paper we have developed a probabilistic approach to the qualitative analysis of the network dynamics of physicochemical models. It is designed to incorporate all available knowledge of the reaction topology, and the parameters on that topology, and calculate the likelihood of achieving an outcome of interest. Inferences on network dynamics are then made by repeating this calculation under changing regulatory conditions and various in silico experiments. We tested the method against a model of the extrinsic apoptosis system and produced qualitative results that were consistent with several lines of experimental research. To our knowledge this is the first attempt at a probabilistic analysis of network dynamics for physicochemical models and we believe this method will prove valuable for the large-scale exploration of those dynamics, particularly when parameter knowledge and data are scarce.

## Acknowledgements

This work was supported by the NIH NCI U01CA215845 (CFL), as well as NSF MCB 1411482 (CFL) We thank the Incyte-Vanderbilt Alliance (CFL) for their support of this project. We also thank the Advanced Computing Center for Research and Education (ACCRE) for computational resources and support needed to complete this work. Finally, we extend our thanks to Dr Blake Wilson for his effort in reviewing this work and providing feedback.

## Supporting information

Table S1

Table S2

Table S3

Table S4

Table S5

Table S6

Figures S1-S4

