## Supplementary Figures and Text for "A probabilistic approach to explore signal execution mechanisms with limited experimental data"

This file includes: Figures S1 to S4

Figure S1: Model calibration to existing FRET data for cleaved Bid, exported Smac, and cleaved PARP.
Figure S2: Log expected values for the six decomposed extrinsic apoptosis networks for increasing values of XIAP.
Figure S3: Apoptotic signal flux over ranges of XIAP and Bcl-2.
Text S1: Interpretation of the data in Figure S3.
Figure S4: Expected value plots at increasing levels of live-points (nested sampling population) for the direct caspase pathway, mitochondrial pathway and complete network.

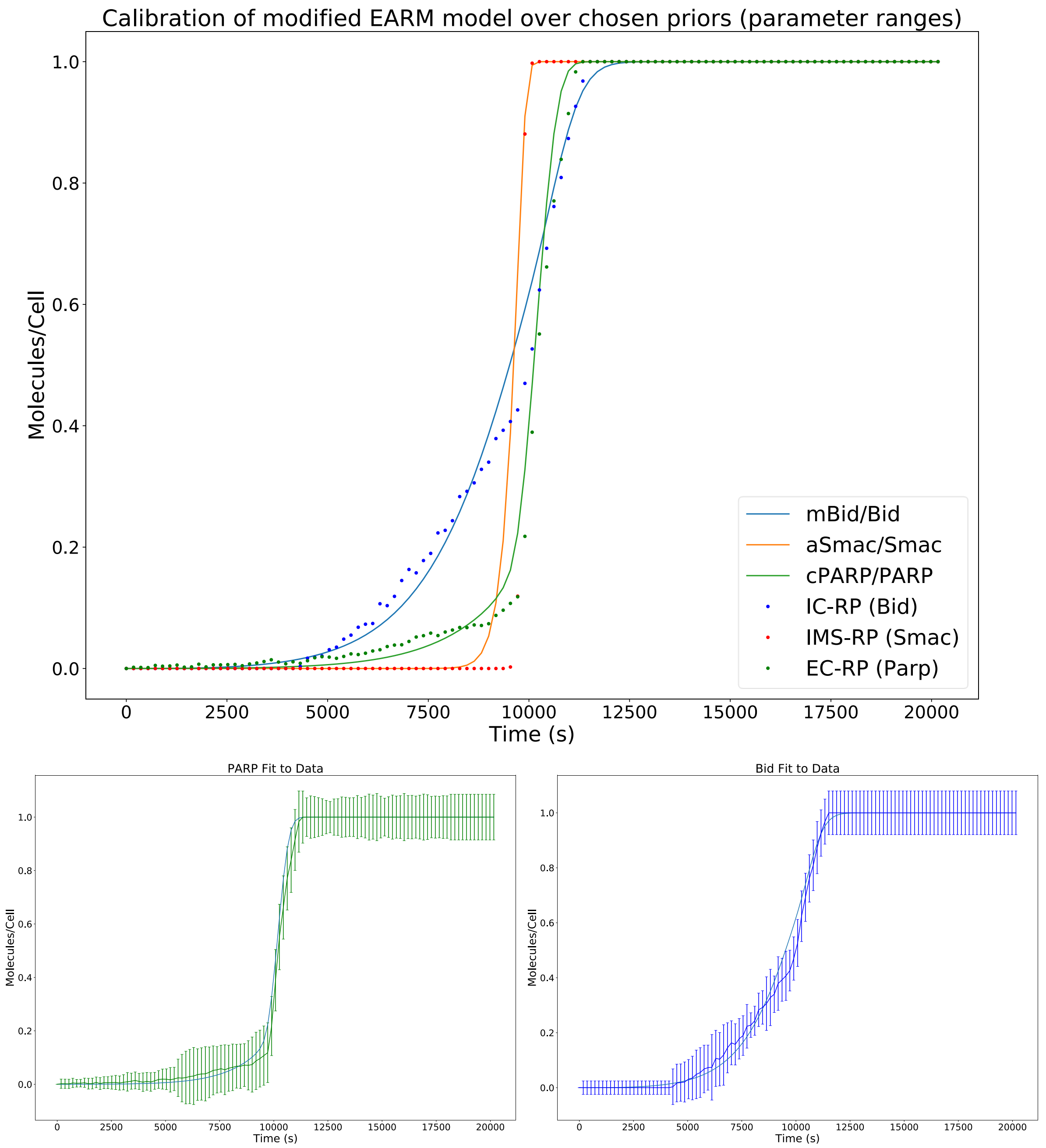


**Figure S1.** (A) Trajectories of cleaved Bid, exported Smac, and cleaved PARP (solid lines) for the calibrated EARM model along with the data used for the calibration (points). (B)-(C) Parp and Bid trajectories respectively along with estimated errors (bars) for the FRET data they were calibrated to.

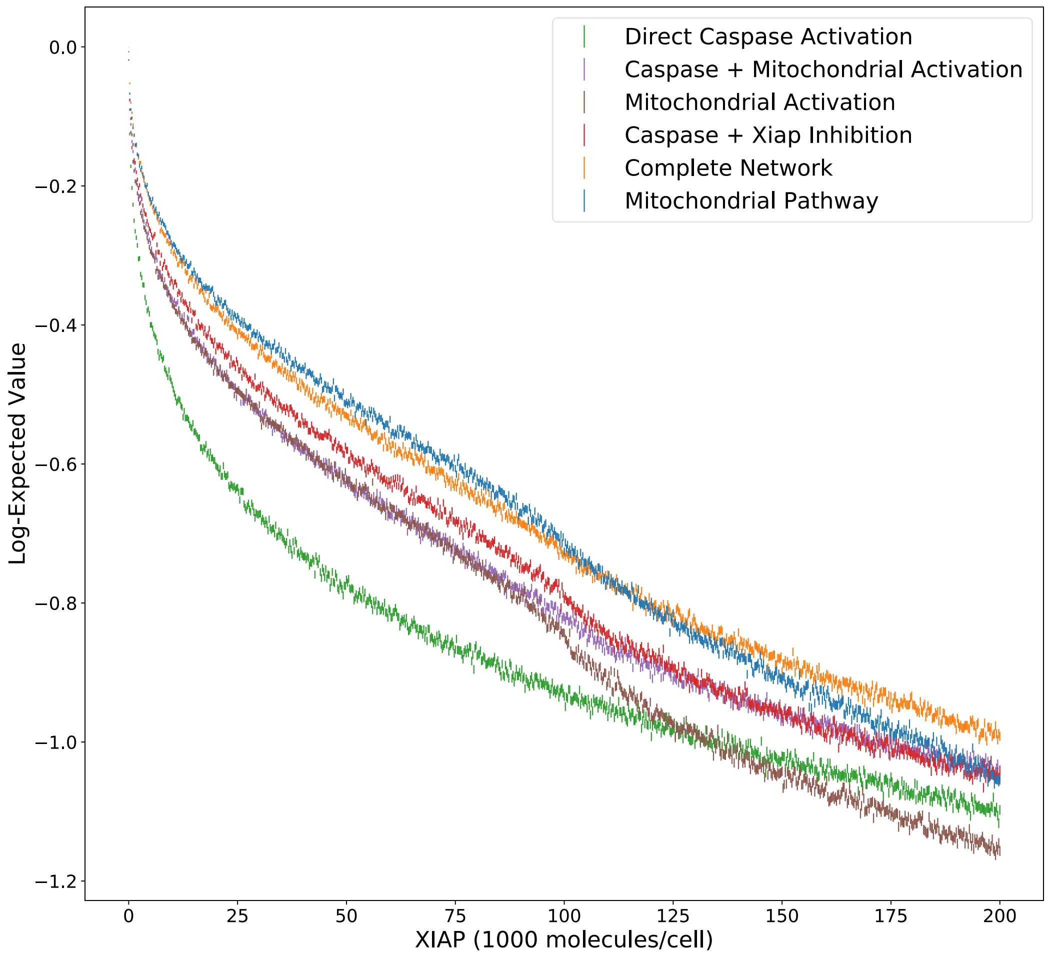


**Figure S2.** log-expected value version of plots in Figure 2G with estimated errors generated by MultiNest.

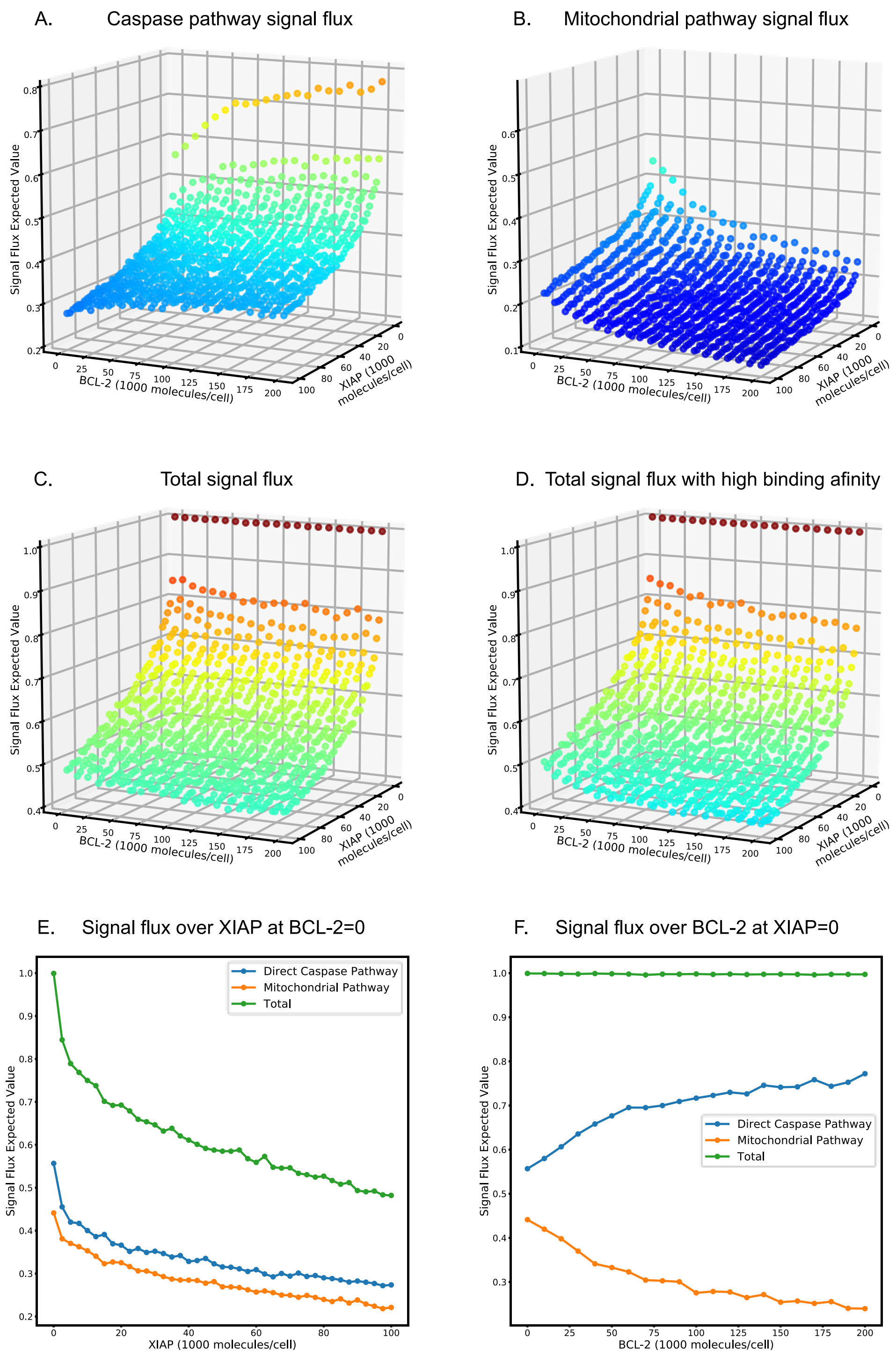


**Figure S3. Apoptotic signal flux over ranges of XIAP and Bcl-2.** (A)-(C) Apoptotic signal flux expected value plots over the apoptosis regulators XIAP (range of 0 to 100,000 molecules per cell) and Bcl-2 (range of 0 to 200,000) for (A) the caspase pathway, (B) the mitochondrial pathway, (C) the total signal flux, and (D) the total signal flux with binding affinities for Bcl-2 and Bad in physiological ranges. (E) and (F) Comparison of the expected value trends at Bcl-2 and XIAP levels of 0 (E and F respectively).

**Text S1. Interpretation of the results from Figure S4.**

Deconstruction of a network into all relevant subnetworks and comparison of the relative changes in apoptosis signal transduction as regulatory conditions change provides a reductionist view of how various network components interact with one another and affect the overall signaling dynamics. To get a holistic view of changes in signaling dynamics that incorporates every network component we calculate the expected values for signal flux through both the caspase and mitochondrial pathways while retaining the complete network model. Instead of considering different models with the same objective function this method considers the same network but equivalent objective functions for different pathway target. Inference of differential signal flow via calculation of pathway flux was introduced in Shockley et al. [1]. In that work they calculated the path fluxes for a number of initial conditions and over an ensemble of parameter sets. Nested sampling, along with a signal flux-based objective function, extends this idea with the calculation of the expected value for signal flux through a target pathway via integration over a prior parameter range. Because both pathways cleave Caspase-3, which goes on to cleave the final product PARP, the objective for each pathway was the sum, over time *T* in seconds, of the proportion of Caspase-3 cleaved through that pathway at time *t* multiplied by the amount of PARP cleaved from time *t-1* to time *t* (see Methods). In addition to runs for both pathways an additional run for the total signal flux was carried out. We explored an XIAP concentration ranging from 0 to 100,000 molecules per cell and Bcl-2 from 0 to 200,000 molecules per cell, in increments of 2500 and 10,000 respectively, producing a 3-dimensional expected value landscape for each target objective. Because the computational cost when using these objective functions is significantly higher than simply calculating the proportion of cleaved PARP at the end of a simulation, the number of live points in the sampling algorithm was reduced from 16,000 to 4,000.

The expected value for signal flux through the caspase pathway showed a sharp initial decline that becomes more gradual as XIAP increases (Figure S4A). This appears to be more pronounced at the higher end of the Bcl-2 range. As Bcl-2 is increased the expected values increase, again becoming more gradual at higher levels of Bcl-2. This effect is clearly more prominent for lower levels of XIAP, likely because of the overall higher levels of signal throughput. The increase in signal flux through the caspase pathway as Bcl-2 increases is indicative of a shifting apoptotic signal from the mitochondrial to the caspase pathway; this is also evident in the decreasing mitochondrial signal flux as Bcl-2 increases (Figure S4B). As with the caspase pathway, increasing XIAP levels causes a decrease in signal flux through the mitochondrial pathway. The effects of both Bcl-2 and XIAP appear to diminish as the other increases. The combined responses to increases in XIAP and Bcl-2 for the caspase and mitochondrial pathways are evident in the expected value landscape for total flux. The total apoptotic signal flux decreases sharply as XIAP increases but remains largely stable, with only relatively small declines, as Bcl-2 is increased. Thus, while XIAP inhibits the flux through both pathways, Bcl-2, under the given simulation conditions, appears to primarily shift the flux to the caspase pathway and inhibits apoptosis to only a small degree compared to XIAP. Note that the expected values for signal flux through the caspase and mitochondrial pathways are additive. The average and average absolute differences between the total flux and the combined caspase and mitochondrial flux are -0.00565 and 0.00866 respectively (Table S4).

Figure S4E displays the expected value trends for each objective over the full range of XIAP and at a Bcl-2 level of 0 for a fully active mitochondrial pathway. Throughout the range of XIAP (and Bcl-2 as well, Figures S4A and S4B), the caspase pathway retains a consistently higher signal flux. This supports the hypothesis that a significant proportion of the mitochondrial signal amplification is due to facilitation of the signal through the caspase pathway via XIAP inhibition and may in fact be the primary mechanism. Figure S4F displays the expected value trends over the range of Bcl-2 at an XIAP level of 0, which clearly shows a shift in flux from the mitochondrial to the caspase pathway. Total flux is nearly complete throughout the range, meaning that the signal results in the cleavage of nearly all available PARP. The shift in signal flux from the mitochondrial to the caspase pathway appears to be, at every level of XIAP and under these simulation conditions, the primary effect of Bcl-2. This implies that even a weak signal through the mitochondria may be enough to inhibit XIAP via SMAC and that much higher binding affinities than the given parameter ranges allowed would be required for Bcl-2 to have a substantial inhibitory effect on apoptosis. Shifting the ranges for the rates of Bcl-2:Bid and Bcl-2:Bax dissociation to [-8.0, -4.0] and [-7.0, -3.0] respectively brings the $K_d$ values roughly in line with [2] and does indeed produce a more pronounced, but still modest, effect (Figure S4D). Further adjustments of the model, both in the initial values and parameter ranges, may be necessary to elicit a higher impact from Bcl-2.

1. Shockley EM, Rouzer CA, Marnett LJ, Deeds EJ, Lopez CF. Signal integration and information transfer in an allosterically regulated network. npj Syst Biol Appl. 2019 Jul 18;5(1):1–9.

2. Ku B, Liang C, Jung JU, Oh B-H. Evidence that inhibition of BAX activation by BCL-2 involves its tight and preferential interaction with the BH3 domain of BAX. Cell Res. 2011 Apr;21(4):627–41.

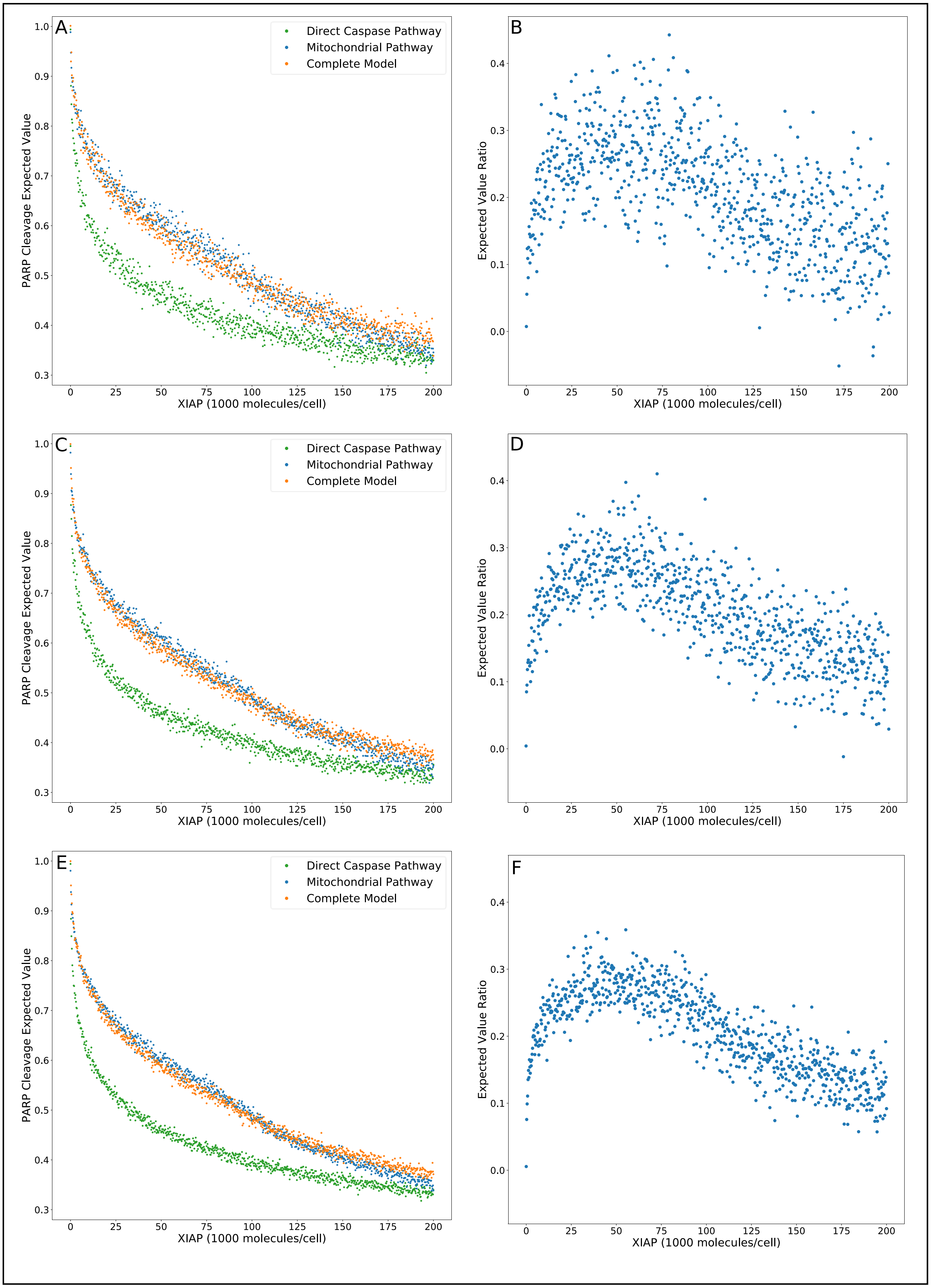
A
B
C
D
E
F
Direct Caspase Pathway
Mitochondrial Pathway
Complete Model
PARP Cleavage Expected Value
Expected Value Ratio
XIAP (1000 molecules/cell)
0
25
50
75
100
125
150
175
200
0.3
0.4
0.5
0.6
0.7
0.8
0.9
1.0
0.0
0.1
0.2

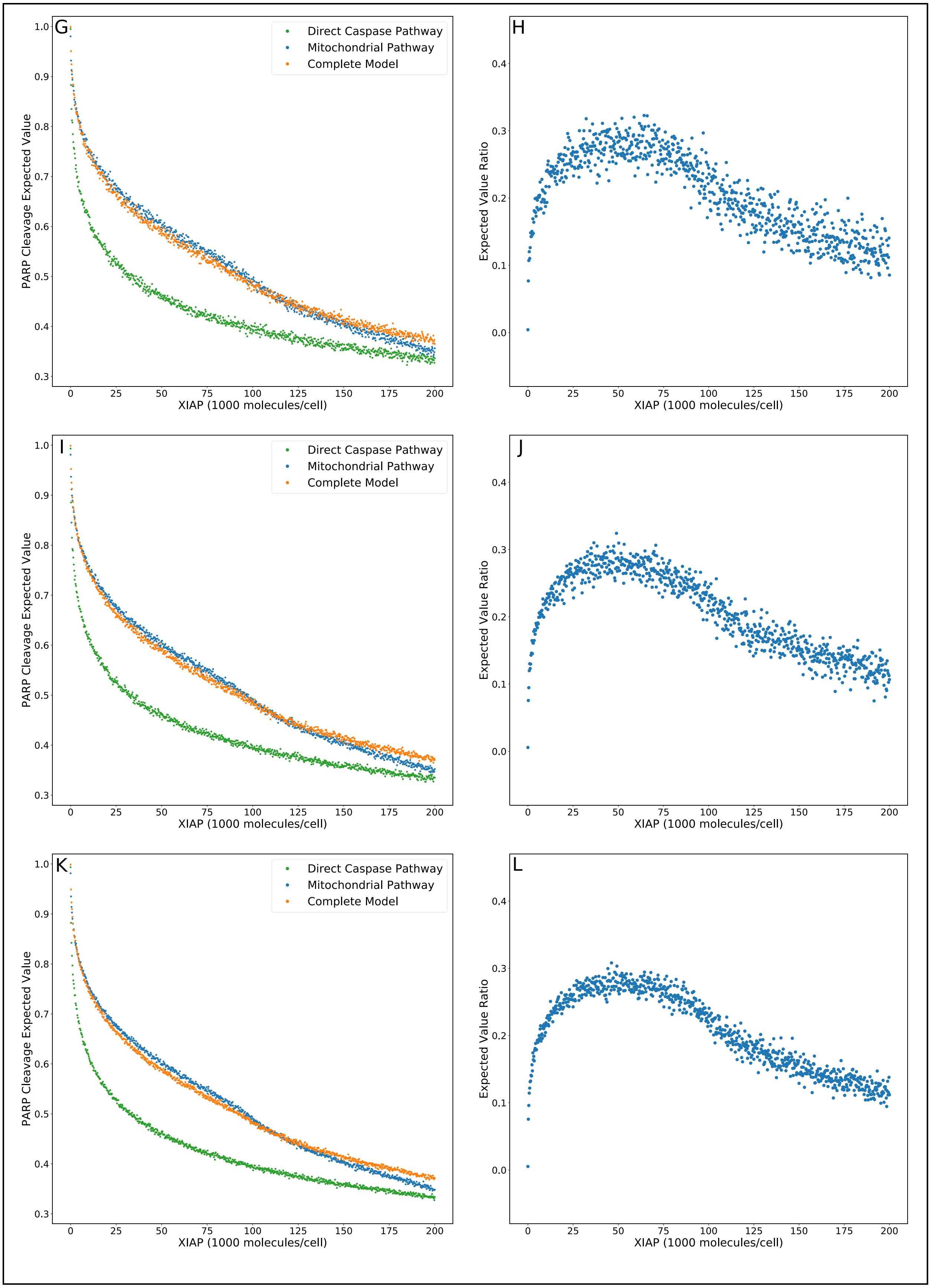
G
H
I
J
K
L
Direct Caspase Pathway
Mitochondrial Pathway
Complete Model
PARP Cleavage Expected Value
Expected Value Ratio
XIAP (1000 molecules/cell)
0
25
50
75
100
125
150
175
200
0.3
0.4
0.5
0.6
0.7
0.8
0.9
1.0
0.0
0.1
0.2
0.3
0.4

**Figure S4.** (A), (C), (E), (G), (I), and (K), expected value plots over increasing levels of XIAP for the direct caspase (green), mitochondrial (blue), and complete (orange) networks with nested sampling population levels of 500, 1000, 2000, 4000, 8000, and 16,000 respectively. (B), (D), (F), (H), (J), and (L), complete/caspase expected value ratio plots derived from the respective expected value plots in (A), (C), (E), (G), (I), and (K).
